# Nanopore adaptive sequencing for mixed samples, whole exome capture and targeted panels

**DOI:** 10.1101/2020.02.03.926956

**Authors:** Alexander Payne, Nadine Holmes, Thomas Clarke, Rory Munro, Bisrat Debebe, Matthew Loose

## Abstract

Nanopore sequencers enable selective sequencing of single molecules in real time by individually reversing the voltage across specific nanopores. Thus DNA molecules can be rejected and replaced with new molecules enabling targeted sequencing to enrich, deplete or achieve specific coverage in a set of reads to address a biological question. We previously demonstrated this method worked using dynamic time warping mapping signal to reference, but required significant compute and did not scale to gigabase references. Using direct base calling with GPU we can now scale to gigabase references. We enrich for specific chromosomes mapping against the human genome and we develop pipelines enriching low abundance organisms from mixed populations without prior knowledge of sample composition. Finally, we enrich panels including 25,600 exon targets from 10,000 human genes and 717 genes implicated in cancer. Using this approach we identify PML-RARA fusions in the NB4 cell line in under 15 hours sequencing. These methods can be used to efficiently screen any target panel of genes without specialised sample preparation using a single computer and suitably powerful GPU.

## Introduction

Selective sequencing, or “Read Until”, is the ability of a nanopore sequencer to reject individual molecules whilst they are being sequenced. We first demonstrated this using dynamic time warping (DTW) against a synthetic reference and selecting for specific regions of a viral genome ^1^. This approach required significant compute resource even though DTW can make use of a number of optimisations ^2^. Others have explored using direct base calling of read chunks to enable Read Until to work direct from sequence ^3^. However, these methods demonstrated little benefit when compared with results obtained without intervention. A promising method currently under development in squiggle space is Uncalled ^4^. Results presented around this method suggests it to have a lighter computational footprint than previous signal based methods, but still requires the use of additional computer resources.

Ideally we would like to work in sequence space as demonstrated by Edwards and colleagues, but also within a reasonable compute framework ^3^. Oxford Nanopore Technologies (ONT) developed a number of base callers for nanopore sequence data, initially utilising Hidden Markov Models and available through the metrichor cloud service ^5^. Subsequently, neural network models were developed which could be run on CPU and then Graphical Processing Units (GPU). To provide real time basecalling, ONT have developed a range of computational platforms with integrated GPU including the minIT, Mk1C, GridION and PromethION ^6^. These devices can provide real time base calling able to keep pace with multiple flowcells generating data. Most recently, these base callers implemented a server-client configuration, such that raw reads containing signal can be passed to the server and a base called sequence returned to the client. Here we show that this GPU base calling can be used to deliver a real time stream of nucleotide data from flowcells sequencing from up to 512 channels simultaneously. At the same time, the GPU can base call the completed reads, enabling dynamic updating of the experiment as results change.

Our method does not rely on comparison of raw current and so we do not have to convert references into signal space as we did for DTW based approaches ^1^. Highly optimised tools such as minimap2 ^7^ can therefore be used to map reads as they are generated and so we can easily update or switch references during sequencing. In this way, selective sequencing can become adaptive sequencing as the sequencer can change its behaviour in response to the data being generated. Our method is only constrained by access to a sufficiently powerful GPU. The results presented here all utilise the ONT GridION MK-1 device which includes an NVIDIA GV100 GPU.

To illustrate the potential of direct GPU base calling for Read Until, we test a range of model problems. Firstly, we select for specific chromosomes from the human genome illustrating that gigabase sized references are not a constraint. Secondly, we investigate the enrichment of low abundance genomes from a mixed population and find we can improve both time-to-answer and the ability to assemble poorly represented genomes. To illustrate adaptive sequencing, we provide a model workflow using centrifuge to identify the most abundant species present within a metagenomic sample, monitor depth of coverage for each in real time and so enrich for the least abundant genomes without *a priori* knowledge of content^8^. This method is necessarily limited by the composition of the reference database and also requires network access to retrieve references once identified. Finally, we enrich panels of genes including 25,000 target regions corresponding to approximately 10,000 genes from the human genome and 717 genes from the COSMIC (Catalog of Somatic Mutations in Cancer) panel ^9^. We demonstrate how Read Until can be used to capture information on key targets without the need for custom library preparation and show we can identify a known translocation in the NB4 cell line in less than 15 hours ^10^. We provide a configurable toolkit which enables targeted sequencing of gigabase genomes including depletion of host sequences as well as example methods to ensure minimum coverage depth for genomes present within a mixed population. Configuration of these tools is relatively straightforward and requires no additional compute as long as sufficiently powerful GPU is available. Our method currently requires that users have access to the Read Until API (ONT).

## Results

### Methods Overview

Selective sequencing requires bidirectional communication with a nanopore sequencer through an API (currently version 2, available from ONT). The API provides a stream of raw current samples from every sequencing pore on the flowcell. Previous API implementations served any signal seen by the Nanopore as a potential read and so required users to process many signals that may not be derived from a genuine read. This caused significant analysis challenges in previous implementations as described by Edwards et al ^3^. The current API version is able to discriminate true DNA signal from background noise more efficiently and so can be configured to only provide data for reads identified as DNA. This reduces the analysis burden on the client side significantly. We reasoned that we ought to be able to convert the signal served by the API into a format compatible with the Guppy basecaller and so retrieve short sequences that could be processed in base space.

Figure S1 illustrates the general workflow for basecalling reads as they are being sequenced. In brief, chunks of read data are grabbed from the Read Until API. The size of these chunks can be configured by the user with the default of 1 second representing approximately 450 bases. Although we have not exhaustively tested all possible values, we found that a chunk size of 0.4 seconds (see methods for configuration information) balanced our desire for the smallest possible chunk size for fast analysis with the possibility of overloading the API with requests for data. This chunk of data, which in theory may contain as many as 512 reads for a MinION flowcell, is processed in one batch by converting the raw signal to Guppy compatible reads and base calling using python bindings to Guppy (PyGuppy, ONT). The returned base called data can then be mapped to a reference using any suitable mapper. Here we use the python API for minimap2, mappy ^7^. Any given read may uniquely map to a specific location in the reference, map to multiple locations, or may simply not map at all. In response the user can choose to reject a read (unblock), acquire more data for that read (proceed) or stop receiving data for the remainder of that read (stop receiving).

We note that 0.4 seconds of signal equates to approximately 180 bases of sequence data (at 450 bases per second) and that the start of a read contains adapter sequence as well as optional barcodes. Read starts can also include some delays as the DNA engages with the pore resulting in a stall signal before signal containing sequence data are available. Thus the first chunk of data may not provide the optimal base call and additional data may be required. However, calling any single chunk of data in isolation is presumably less informative than calling the entire signal. We therefore implement a read cache which concatenates adjacent signal data from the same read as it is acquired from the API. This cache can keep up with the 0.4 second API queries. This enables calling the complete available signal for each read since it began. In a typical experiment we find that 90% of reads can be called and mapped within 3 chunks (1.2 seconds) using this method (Figure S2).

The ONT Guppy basecaller contains at least three models for basecalling. These models trade speed (fast) for accuracy (hac) and can optionally call methylation. For selective sequencing, the goal would appear to be speed and so we investigated the efficacy of selective sequencing using both the fast and hac models. However, we found that the high accuracy model (hac) was able to keep up with the rate of data generation on the GridION Mk1. Across all the experiments presented in this paper, the average time to call a single batch of reads was 0.28s and the average batch contained 30 reads. Thus we can call at least 100 read chunks per second (see Figures S3-7).

Depending on the experiment that a user is seeking to perform, the response to a read mapping to a specific sequence in a reference genome may vary (see online methods). Clearly, if a user is trying to deplete reads originating from a host then a read mapping to that host should be rejected. Likewise, when seeking to enrich for a target, reads mapping to that target should be sequenced. Less well considered are reads which do not map to a reference at all. If the goal of the experiment is to enrich low abundance or unknown targets, these reads should be sequenced. If enriching for subsets of a known reference, these reads might be best rejected in favour of sampling more molecules. Given the variety of options, we developed a configuration file which allows any mapping results to be mapped to any action. We have also included the option to dynamically update a TOML configuration file during a sequencing run, enabling targets to be changed on the fly. Optionally the TOML file can be configured to carry out different experiments on different regions of the same flowcell (see https://github.com/LooseLab/ru/blob/master/TOML.md).

## Read Until Performance

### Enrichment and Depletion

To test the performance of our real time base calling approach, we sequenced the well studied NA12878 reference cell line ^11^. We configured the flowcell to operate in four separate quadrants. The first acted as a control (all reads accepted), the second sequenced reads derived from chromosomes 1-8 (50% of reads accepted), the third sequenced chromosomes 9-14 (25% of reads accepted) and the fourth chromosomes 16-20 (12.5% of reads accepted). The only differences between quadrants are the targets for rejection: all reads are basecalled and mapped to the reference regardless of origin. The median read lengths per chromosome in each quadrant indicate which are being sequenced and which are being rejected (Figure 1A). Selectively sequenced reads have a median read length of approximately 15 kb. In contrast, rejected reads have a median length of approximately 500 bases, equating to approximately 1.1 seconds of sequencing time. Thus reads have been basecalled, mapped and the unblock action sent and actioned within approximately 1 second of the read starting. This run yielded 9.5 Gb of sequence data, but was not evenly distributed across the four quadrants. 3.47 Gb were generated in the control portion, 2.79 Gb in the 50% acceptance, 1.84 in the 25% acceptance and only 1.22 Gb in the 12%. This drop in efficiency can be seen in a heatmap of bases sequenced per channel over the course of the experiment (Figure 1B). For the quadrants, the optimal enrichment is 2-fold, 4-fold and 8-fold but we see lower enrichments by the end of the experiment, presumably due to the lower yield (Figure 1C). Importantly, we do see enrichment of our target sequences in all cases compared with the control region of the flowcell. Notably, relative enrichment is greater at the beginning of the sequencing run and is closer to the theoretical maximum enrichment we expect to see (Figure 1D). Analysis of the number of channels contributing to data generation over the course of the run shows that sequencing capacity is lost faster as more reads are rejected (Figure 1E). Although we did not use nuclease flush on this run we would anticipate improvements, in both yield and enrichment, with this method. Performance metrics for Read Until in this experiment are shown in Figure S3 and we were able to call all batches within our 0.4 second window.

**Figure 1.**
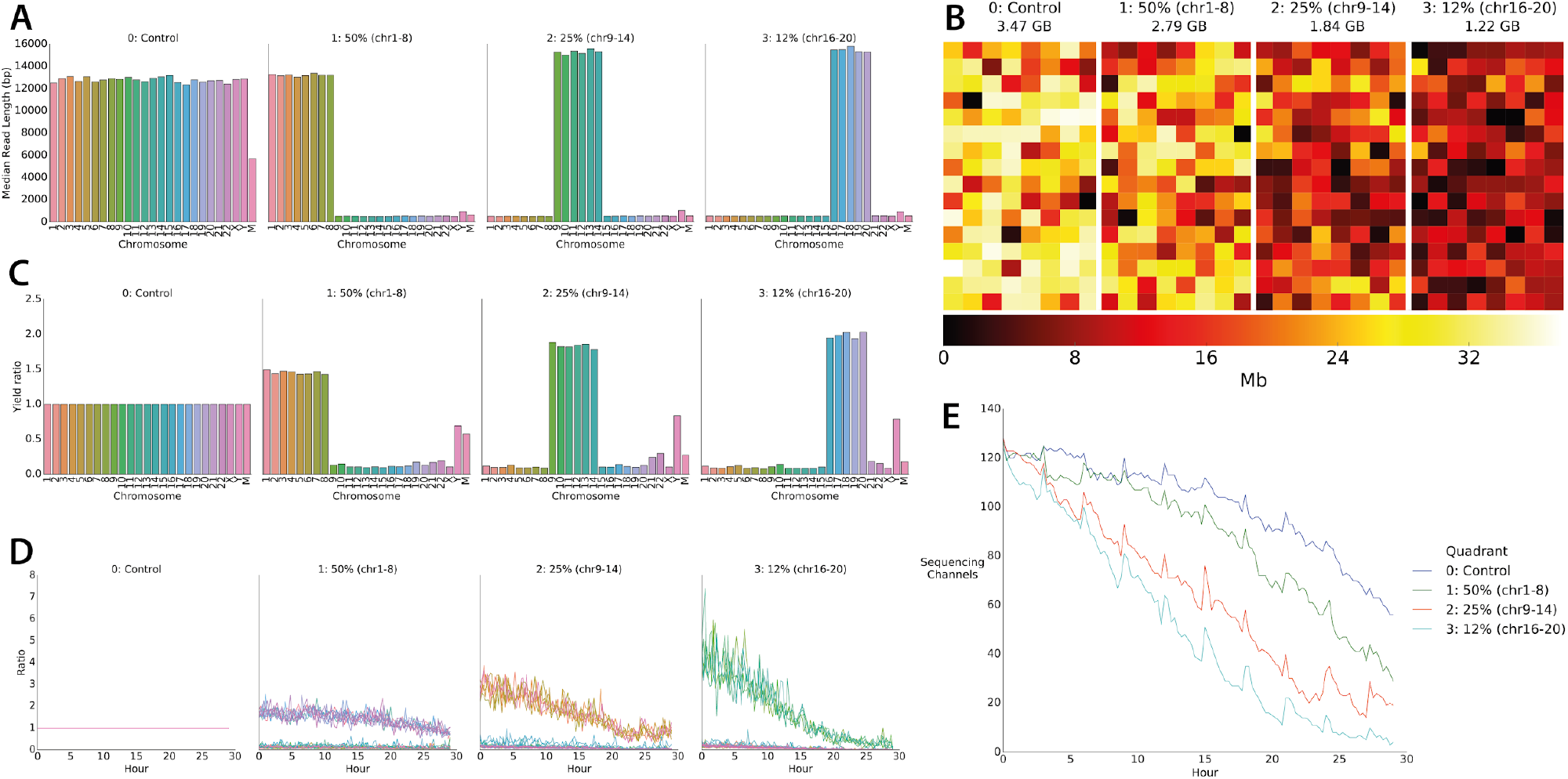
Human Genome Scale Selective Sequencing. A) Median read lengths for reads sequenced from GM12878 and mapped against HG38 excluding alt chromosomes. The four panels each represent a quadrant of the flowcell. In the control all reads are sequenced, in the second reads mapping to chromosomes 1-8, in the third reads mapping to chromosomes 9-14 and the fourth reads mapping to chromosome 16-20. The combined length of each of these target sets equates to approximately ½, ¼ and ⅛ of the human genome respectively. B) Heatmap of throughput per channel in each quadrant from the flowcell illustrating reduced yield as the proportion of reads rejected is increased. C) Yield ratio for each chromosome in each condition normalised against yield observed for each chromosome in the control quadrant. D) Yield of on target reads calculated in a rolling window over the course of the sequencing run showing the loss of enrichment potential. E) Plot of the number of channels contributing sequence data over the course of the sequencing run. Channels are lost at a greater rate when more reads are rejected.

### Enrichment of metagenomes and “Run Until”

A common goal in sequencing library preparation is to remove host DNA and so enrich for a subpopulation of material ^12,13^. This is a common application for which selective sequencing might be beneficial, although most likely in conjunction with library preparation methods. We therefore considered metagenomics applications as a similar class of problem. Nicholls et al demonstrated reference sequencing of a mock microbial standard across both GridION and PromethION flowcells ^14^. Using a mock community available from ZymoBIOMICS they were able to generate sufficient data to assemble several of the bacteria into single contigs without an intermediate binning step. Notably the eukaryotic genomes which were present within the sample at lower abundance (2%) did not generate high contiguity assemblies. This is not particularly surprising as the coverage depth for *Saccharomyces cerevisiae* was only 17x and *Cryptococcus neoformans* only reached 10x with data from a single GridION flowcell ^14^. Enriching for these low abundance components is conceptually similar to depleting host material from a sample.

To maximise the benefit of selective sequencing, we used the ZymoBIOMICS high molecular weight DNA sample (ZymoBIOMICS). The use of this sample will *a priori* improve assemblies due to the longer read lengths obtained. This sample also differs from that used by Nicholls et al. as it does not include *Cryptococcus neoformans. Saccharomyces cerevisiae* is included at approximately 2%. To see if selective sequencing could improve the relative coverage of low abundance material we developed a simple pipeline (ru_iteralign) to drive our selective sequencing decisions (Figure S1). In brief, ru_iteralign maps base called reads that have been completely sequenced against a reference as they are generated. Once the depth of coverage for a particular reference sequence reaches a pre-specified mean coverage level, the sequence can be dynamically added to the Read Until TOML configuration file causing further reads mapping to this sequence to be rejected. Here, we rely on the GPU built into the GridION mk1 to simultaneously process both the real time data stream for Read Until and the normal base calling activity of the device. We utilised the MinKNOW API to log the point at which each reference reached the specified target coverage in the MinKNOW interface. We also implemented Run Until to stop the run automatically once all targets had reached a predetermined sufficient coverage.

These experiments use a specific reference file for this community. Mean read lengths for each target genome reduce once the specific target is added to the rejection list (Fig 2A), as the mean read length becomes dominated by short, rejected reads. Plotting coverage over time for reads that were not rejected by Read Until shows a corresponding decrease in coverage accumulation for genomes which have reached the desired coverage level with an increase in sequencing potential for the least abundant sample, *Saccharomyces cerevisiae* (Fig 2B). Visualising the proportion of bases mapping to each genome over the course of the run reveals the shift in sequencing capacity to *Saccharomyces cerevisiae* (Fig 2C). Illustrating that relative abundance in the sample can still be determined when running Read Until, the relative proportions of reads mapping to each genome does not change whilst reads are being rejected (Fig 2D). We configured this run to automatically stop once each genome represented in the ZymoBIOMICS sample reached a coverage of 40x, which took approximately 16 hours (performance metrics in Figure S4, this run used an earlier version of Guppy and was less performant).

**Figure 2.**
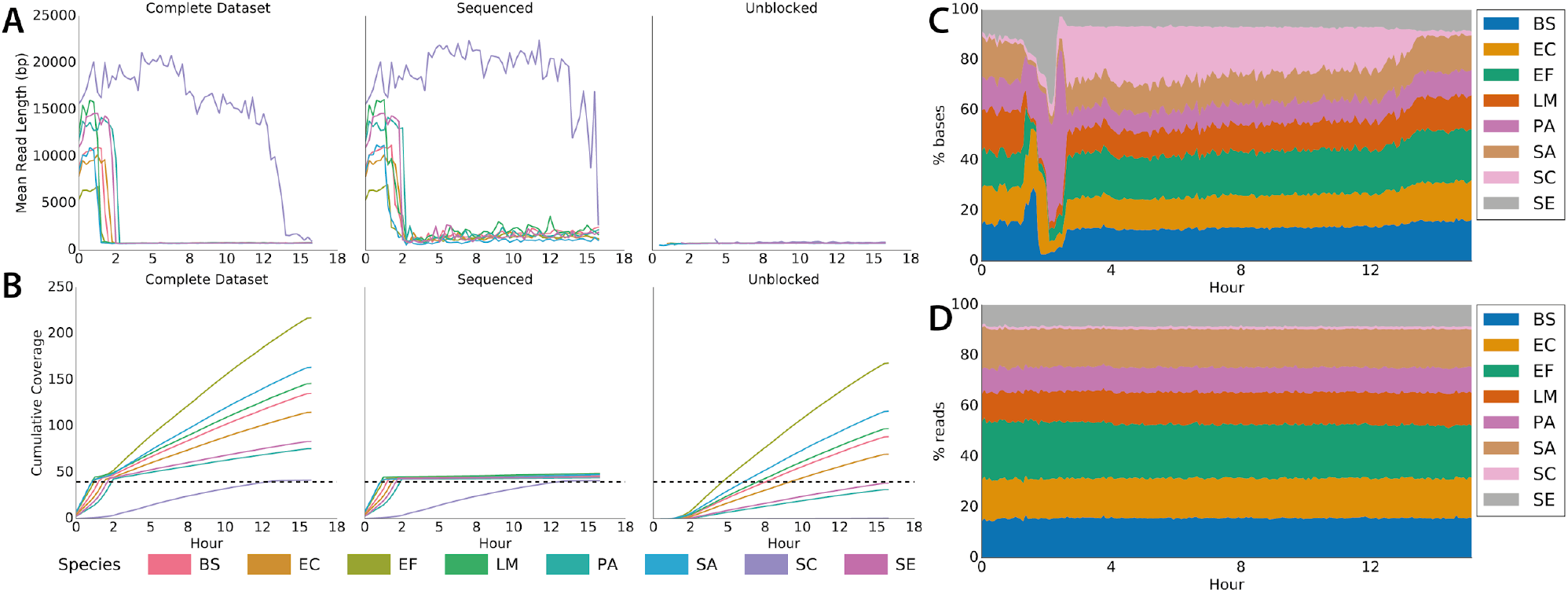
Adaptive sequencing enriching for the least abundant genome and ensuring uniform 40x coverage. A) Mean read lengths for reads sequenced from the ZymoBIOMICS mock metagenomic community mapped against the provided references (ZymoBIOMICS, USA). Read lengths are reported for the whole run, the deliberately sequenced reads and those which were actively unblocked. B) Shows cumulative coverage of each ZymoBIOMICS genome during the sequencing run. The total coverage still accumulated as unblocked reads, though short, still map. Sequencing was automatically terminated once each sample reached 40x. C) Stacked area graph illustrating how the proportion of bases mapping to each species changes over time. D) In contrast, the proportion of reads mapping to each species over time doesn’t change significantly.

In order to reach 40x, this run generated 4.4 Gb of sequence data. According to ZymoBIOMICS, this sample should be 2% *Saccharomyces cerevisiae* by bases. This would yield approximately 88 Mb of sequence data for *S. cerevisiae* or approximately 7x coverage. Using selective sequencing we obtained 40x coverage for *S. cerevisiae*. Naively this represents a 5.7 fold increase in on target data generation. However, given a flowcell not implementing selective sequencing would likely have higher yield, real world enrichment is lower. For example, Nicholls et al report 16 Gb on a similar ZymoBIOMICS run ^14,15^. Similar yields here would have resulted in approximately 25x coverage of *S. cerevisiae* giving an effective enrichment of 1.6x. In theory, enrichment of a 2% subset of a sample should be greater than this, but as we saw with the human experiments above, there is a cost to rejecting an individual read. Even so, we find in multiple experiments (n=3) we could enrich the least abundant element compared with that expected from the sample composition. Although the enrichment over the whole run is low, we find we accelerate the time-to-answer at which a particular coverage depth can be achieved.

This approach assumes knowledge of the composition of the sample *a priori* and so is of limited practical relevance. Therefore we integrated a metagenomics classifier into our pipeline (ru_iteralign_centrifuge) ^8^. We chose centrifuge for its low memory footprint and the straightforward way in which it can be used iteratively. In this approach, we do not map reads to a reference until they have been classified by centrifuge. We track the numbers of reads mapping to specific taxonomic IDs and identify those that pass a user defined threshold (here set to 2000). Appropriate reference genomes corresponding to those identified by centrifuge are retrieved from RefSeq^16^ and subsequent reads are both classified with centrifuge and mapped against the expanding set of references. Once target coverage depth is reached (here set to 50x), the Read Until configuration TOML is updated with the reference sequences and list of targets to reject by updating the TOML files configuring the tool as with ru_iteralign. This illustrates the principle of adaptive sequencing.

This run generated 5.995 Gb of sequence data and all bacterial genomes present within the sample were successfully identified. Rejected read lengths were slightly longer than our previous experiments, but enrichment was still achieved although we did not obtain 50x coverage of *S. Cerevisiae* before the flowcell was completely blocked (Fig 3, Figure S5,S8). 6 Gb of sequence data should result in approximately 10x coverage. However, a run achieving 16 Gb would result in approximately 26x coverage. Again, the benefit here is a combination of time-to-answer and slight enrichment. 50x coverage without selective sequencing would require a 24 Gb sequencing run likely to take 48 hours or more on a single flowcell. The experiment presented here was complete within 24 hours. As expected, improved coverage depth results in more contiguous assemblies using MetaFlye (Fig S9); however this is in part a consequence of the improved read lengths in these assays ^14,17^. It is likely that subsequent nuclease flushing of the flowcell would increase effective throughput, but as our goal was to reach a target coverage within a time frame we did not need to test this.

**Figure 3.**
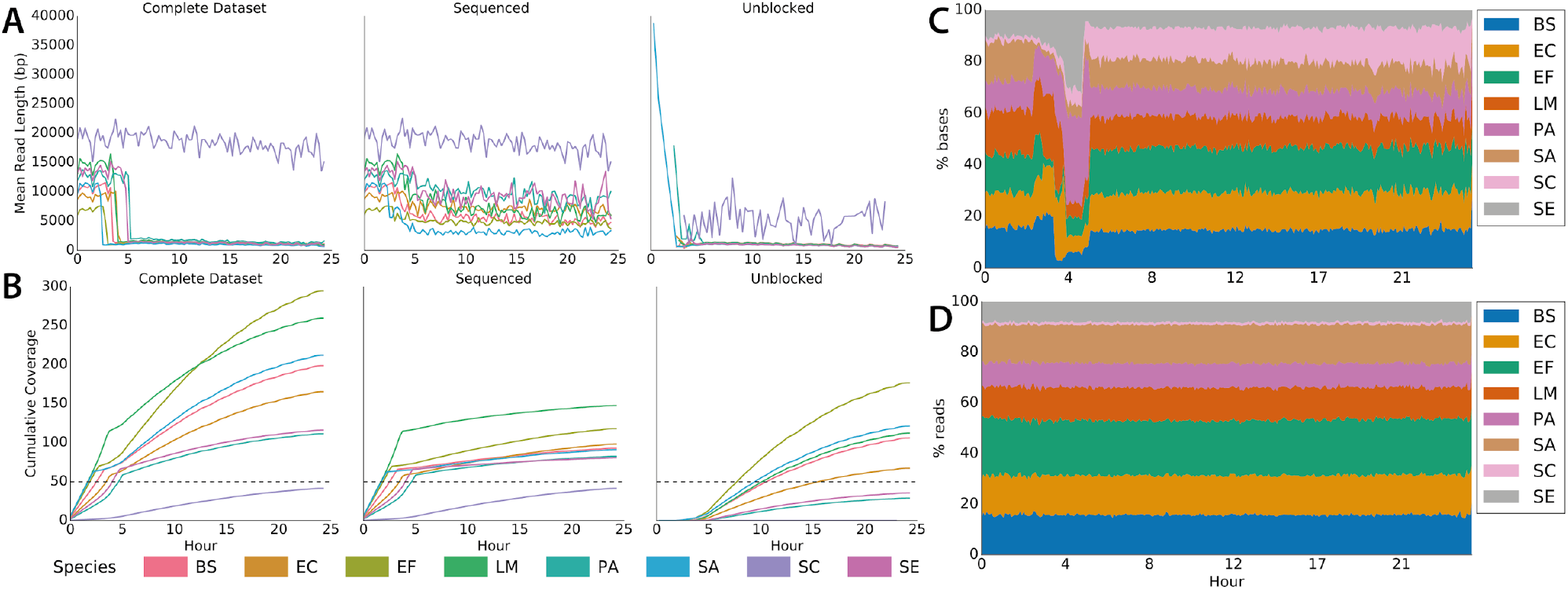
Adaptive sequencing enriching for the least abundant genome with centrifuge read classification and ensuring uniform 50x coverage. A) Mean read lengths for reads sequenced from the ZymoBIOMICS mock metagenomic community mapped against the provided references. Read lengths are reported for the whole run, the deliberately sequenced reads and those which were actively unblocked. B) Shows cumulative coverage of each ZymoBIOMICS genome during the sequencing run. The total coverage still accumulated as unblocked reads, though short, still map. Sequencing was automatically terminated once each sample reached 50x. The small overshoot in sequenced reads coverage is likely caused by the centrifuge step. C) Stacked area graph illustrating how the proportion of bases mapping to each species changes over time. D) In contrast, the proportion of reads mapping to each species over time doesn’t change significantly.

### Target Panel Enrichment

There are many methods available for target enrichment in sequencing panels including PCR amplification and bait capture methods ^18,19^. More recently methods for capturing specific regions of genomes with CRISPR-Cas9 have been used to enrich DNA extractions and libraries prior to sequencing ^20^–^22^. These methods can provide reliable and cost effective screening panels when applied at scale, but have associated development, instrument and consumable costs. Unlike methods which capture native DNA ^20^, PCR based methods cannot be used to capture methylation information without additional processing of input samples. There is also no way of tuning a panel once a sample has been prepared.

Selective sequencing is a tempting alternative and so we sought to capture targets from the human genome. Given we can scale to the human genome, we selected 19,296 target genes annotated as protein coding with Transcript Name IDs from GCRh38 excluding those on alt chromosomes, X and Y ^23,24^. We extracted exon coordinates, extended 3kb either side and collapsed overlapping targets together. We then chose to enrich for only those targets found on odd numbered chromosomes. This gave a total search space of 176 Mb in the human genome (approx 5%) containing 25,600 target regions covering nearly 10,000 genes (Figure 4A). On a single GridION flowcell with 1,660 pores we obtained 6.1 Gb of sequence data in 24 hours. After a nuclease flush, loading additional library and sequencing for a further 24 hours gave an additional 5.573 Gb (total yield: 11.675 Gb, N50 9 kb). We found that our exon targets had mean coverage of 17.39x (median 17.23x) with 75%>14.15x and 25%>20.42x. On the “control” even chromosomes, median coverage of depleted targets was 0.98x (mean 1.2x). Detailed coverage plots of targets on ODD (Figure 4C,D) and EVEN (Figure 4E,F) chromosomes shows how coverage correlates with the target regions used in the experiment. Controlling for these experiments is complicated by flowcell variability. Therefore we compare with theoretical yields of 10, 20 and 30 Gb resulting in approximately 3-10x coverage. On this basis, our effective enrichment is from 2.7-5.4x depending on flowcell performance, consistent with our earlier observations on the human genome. Nuclease flushing significantly assists in this enrichment and the effect of this can be seen in flowcell performance metrics (Figure S6).

**Figure 4.**
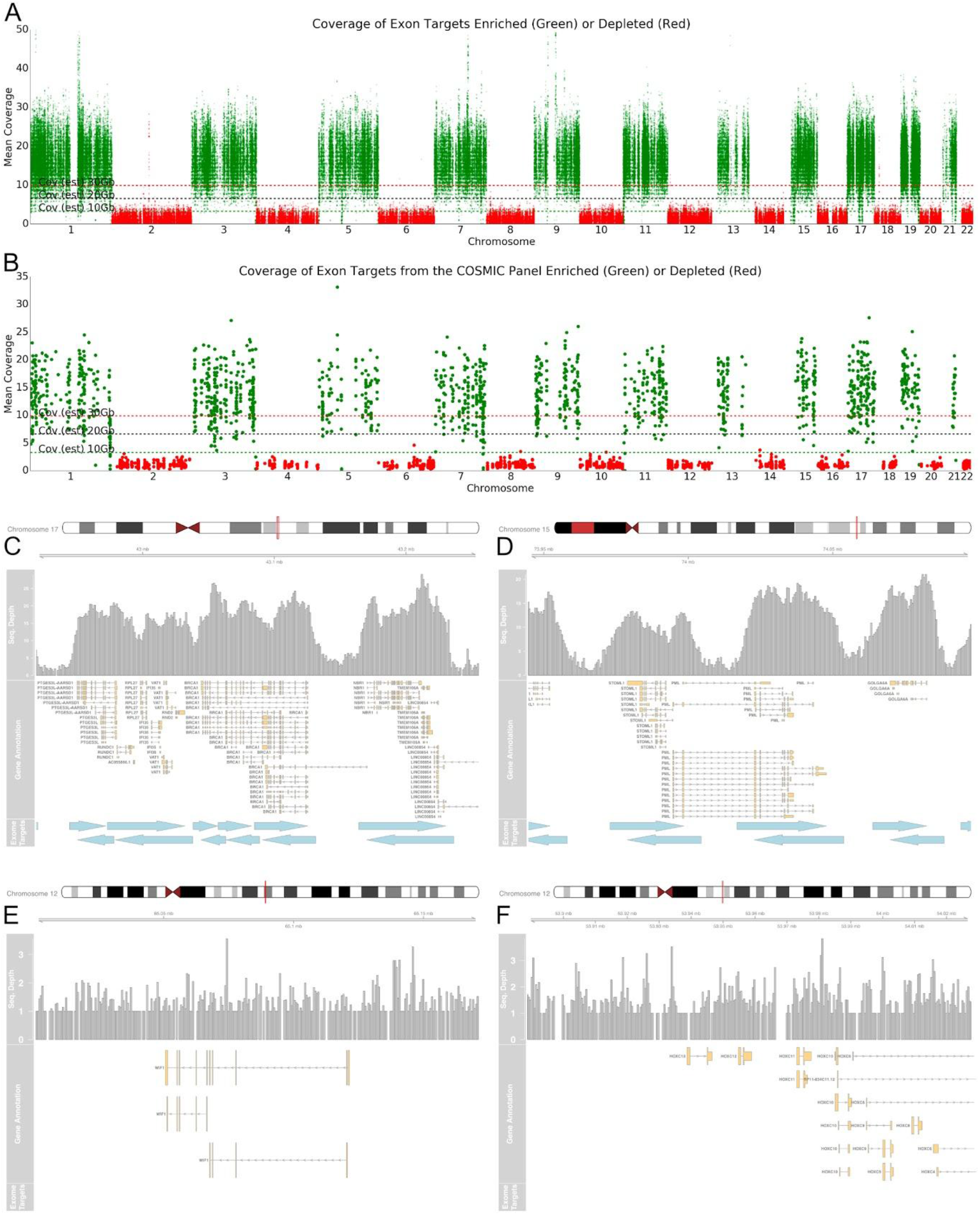
Half Exome Panel Targeted Sequencing. A) Mean coverage across each exon target in the genome ordered by chromosome. Exons on odd numbered chromosomes are enriched (green) and depleted on even numbered chromosomes (red). B) Mean coverage across each exon for genes within the COSMIC cancer panels. Horizontal lines represent approximate mean expected coverage for flowcells yielding 10, 20 or 30 Gb of data in a single run. Mean coverage calculated by mosdepth^25^. C,D,E,F) Coverage plots for highlighted genes including BRCA1 (C), PML (D), WIF1 (E) and HOXC13 and HOXC11 (F). C and D are enriched as they are found on chromosome 17 and 15 whilst E and F are depleted as genes are on chromosome 12. Exon target regions indicated by blue bars. In this experiment, different targets were used for the watson and crick strands as illustrated by the offsets.

Coincidentally, our initial exon panel contains 371 genes from the Catalogue of Somatic Mutations in Cancer (COSMIC) ^9^. Target sequences (exons and surrounding overlaps) had median coverage of 13.7x for these sequences (Figure 4B). Figure 4C and D illustrate the coverage obtained over BRCA1, PML and their surrounding targets. In our exon capture approach, we excluded intronic sequences to reduce the total search space (although this isn’t a requirement of our approach), however it would be preferable to sequence completely through these targets. To illustrate the flexibility of our approach, we therefore generated a target panel covering the entire COSMIC panel (717 genes) excluding those with no genomic coordinates in the COSMIC database (see Supplementary File 1). In total, this covers 82.75 Mb of target sequence including introns. We added an additional flanking 5 kB either side, resulting in a total search space of 89.9 Mb (approx 2.7%). We sequenced this panel using a flowcell with a high pore count (approx 1,724 at the start). The first run generated 3.7 Gb within 24 hours. After nuclease flush and reload a further 6.03 Gb were generated giving a total of 9.73 Gb (Performance metrics in Figure S7). The read N50 was 940 bases reflecting the overall efficiency of read rejection. Deliberately rejected reads had an N50 of approximately 515 bases with the N50 of sequenced reads at 11,564 bases. Over gene target regions the median coverage was 32.2x (mean 30.7x) (Figure 5A, Supplementary File 1), with 75% of genes >28x, 25% of genes > 35x.

**Figure 5.**
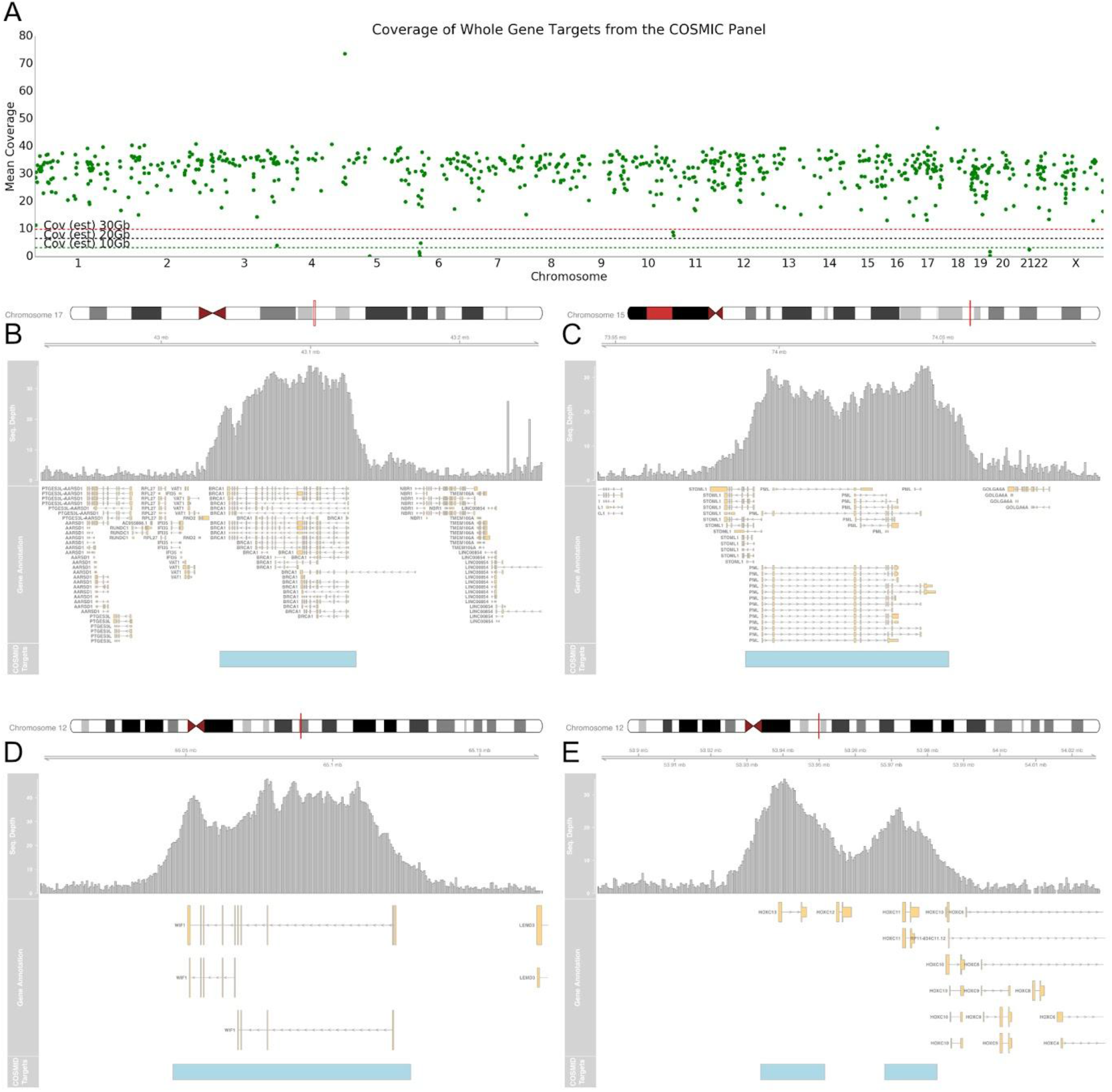
COSMIC Panel Targeted Sequencing. A) Mean coverage across the selected COSMIC gene regions ordered by chromosome. Horizontal lines represent approximate mean expected coverage for flowcells yielding 10, 20 or 30 Gb of data in a single run. Mean coverage calculated by mosdepth^25^. B,C,D,E) Coverage plots for highlighted genes including BRCA1 (B), PML (C), WIF1 (D) and HOXC13 and HOXC11 (E). COSMIC Target regions indicated by blue bars.

Figure 5B-E shows the coverage observed for BRCA1, PML, WIF1 and HOXC13 and HOXC11. The specificity of selective sequencing can be clearly seen, particularly around the Hox genes where neighbouring members of the HOXC cluster are not sequenced. In this example, we did not alter the boundaries for sequencing on the forward and reverse strands meaning that additional gene panels can be generated from a simple BED file. As seen in Bowen et al these data could be used to assess methylation, structural variants and given sufficient depth, nucleotide variation ^20^.

To test if our method can be used to screen for structural variants we used DNA extracted from the NB4 acute promyelocytic leukemia (APL) cell line ^10^. We sequenced this sample using the same COSMIC cancer panel to determine if we could identify the translocation (Figure 6A, Figure S10). Sequencing was carried out on a flowcell with only 1,196 pores and generated 4.5 Gb of sequence data in less than 15 hours. Over gene target regions the median coverage was 11.46x (mean 11.78x) (Figure 5A, Supplementary File 1), with 75% of genes >9.5x, 25% of genes > 13.4x. Analysis of these reads with svim looking for breakpoint ends, and ignoring in/dels, identified just two candidates passing default svim filtering (see methods) ^26^. This breakpoint can also be detected with sniffles (data not shown) ^27^. Of these, one captured the known breakpoint in this sample supported by six reads (compare Figure 6B,C). Subsequent collection of additional data (24 hours of further sequencing generating 3Gb more of sequence) resulted in median coverage of 17.37x (median 18x) with 11 reads supporting the variant. Within the reference NA12878 cell line material no complex rearrangements were reported in genes enriched with the COSMIC panel (Figure 6C). Notably we did observe several potential such events in the wider panel (see Supplementary Table 1).

**Figure 6.**
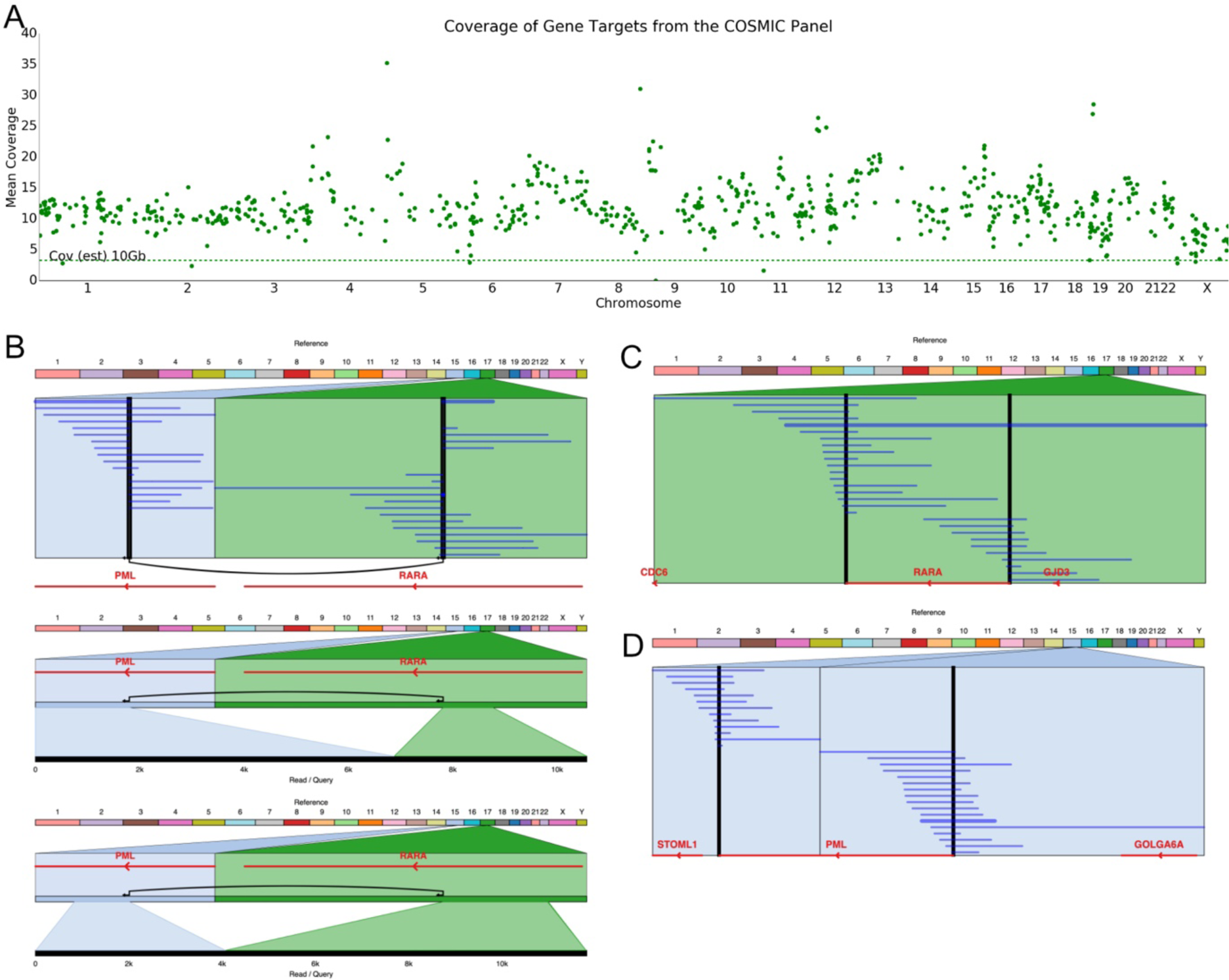
COSMIC Panel Targeted Sequencing of NB4. A) Mean coverage across each of the COSMIC target regions ordered by chromosome. Horizontal dashed line indicates expected coverage from a flowcell yielding 10Gb of sequence data in a single run. B) Reads mapping to chromosomes 15 and 17 derived from the NB4 cell line indicating the fusion between PML and RARA. Mappings of two example individual reads shown below. C) Reads derived from NA12878 covering the breakpoints identified from analysis of the NB4 cell line showing no rearrangement in RARA. D) As C but illustrating PML. Black vertical lines illustrate the coordinates of breakpoint ends as identified in NB4. Breakpoints identified using svim, visualisations using Ribbon ^26,28^.

## Discussion

The idea of selectively sequencing (‘Read Until’) individual molecules using only computational methods is a unique aspect of Nanopore Sequencing ^1^. Here we exploit ONT tools to provide a true real-time stream of sequence data as nucleotide bases. This approach removes the need for complex signal mapping algorithms but does require a sufficiently performant base caller. Prior work by Edwards and colleagues illustrated that this method was a feasible approach, but required extensive additional compute and was unable to show significant enrichment over throughput achieved without running ‘Read Until’ ^3^. Here we demonstrate real enrichment over that expected from a similar control flowcell. We exploit techniques for enhancing flowcell yield such as nuclease flushing and loading additional library, but these same methods would also be required to maximise throughput on a standard sequencing flowcell.

Key benefits of the approach shown here are that we utilise only compute resource available within the GridION Mk1. As we use current commercially provided base callers, we can utilise new algorithms and pores as they are developed. Thus, although not yet tested, we could use this method on RNA if sufficiently long reads require depletion. Similarly we could use methylation aware base callers to sequence regions of DNA from either high or low methylation regions. As we ultimately obtain sequence, rather than signal, data we dramatically simplify the construction of pipelines for downstream analysis of reads. Although we have only shown results for the GridION Mk1 this method should be applicable to any MinION configuration with sufficient GPU to basecall a sequencing run in real time. Early tests on systems configured with NVIDIA 2080 GPUs suggest they can keep up with a single flowcell in real time (J Tyson, Pers. Comm.). In principle this method should scale to the PromethION.

We do find that increased rejection of reads on a flowcell negatively impacts the total sequencing yield and so impacts the enrichment observed. Although we have not extensively exploited methods to wash and reuse flowcells here, where we have tried this we do see increased yield and enrichment. As a consequence, the current main benefit of selective sequencing in metagenomics and host depletion is to improve time-to-answer. For samples which sequence well (i.e do not tend to block the flowcell), additional enrichment benefits may be observed. Notably running selective sequencing does not disrupt the proportion of reads by count that map to a specific reference. Thus in the case of metagenomics type applications it is still possible to assess relative abundance whilst focussing sequencing length on specific subsets of reads. Future methods proposed by ONT to address blocking such as onboard nucleases might increase throughput in future.

We demonstrate that selective sequencing of arbitrary targeted regions of the human genome results in actionable coverage and we can identify structural variants within the COSMIC cancer panel. In total, DNA extraction, library preparation, sequencing and analysis could be completed within 24 hours. When sequencing a subset of a large genome, large numbers of off-target reads are sampled whilst detecting those of interest and the precise parameters of optimal target size and coverage have yet to be defined. As a consequence, library preparation methods that enrich for regions of interest will likely result in higher coverage compared to ‘Read Until’. However, the design of such panels is relatively costly and inflexible once developed. Any method that relies on amplification results in the loss of methylation data which would otherwise be found in these samples. Of course, throughput achievable on platforms such as the PromethION at scale provide whole genome sequencing at relatively low cost^29^. Thus any effective method for enrichment must compete with these costs, including the additional compute required. By utilising the available GPU compute capacity during the sequencing run, we address this issue. There is no reason, in theory, why samples could not be multiplexed on a single flowcell as long as sufficient yield can be obtained to address the biological question.

In selective sequencing, targets can be updated by configuring a single configuration file. Developing a new panel is as straightforward as compiling a list of target regions. Here we also illustrate the concept of adaptive sequencing, as in our metagenomics examples, where targets can be dynamically adjusted during a run. In theory a panel could be updated in response to observations of the data in real time, perhaps adding targets where candidate novel structural variants have been identified or removing targets where sufficient evidence is available to eliminate the possibility of an SV existing.

Although we have focussed exclusively on applications for read until, we believe that a real time sequence data stream in bases has significant advantages for future pipelines. If sequence data can be streamed directly into an analysis pipeline and conclusions drawn without the requirements for data storage then field deployment of sequencing for detection of specific sequences might be accelerated. Ultimately it may be possible to stream sequence data for calling of structural variants and further analysis in real time.

## Online Methods

### Library preparation and sequencing

Standard LSK-109 (ONT) sequencing libraries were prepared from either the ZymoBIOMICS HMW DNA Standard (DS6322 ZymoBIOMICS USA) or DNA extracted from GM12878 cells (Coriell), or NB4 cells (gift from M. Hubank) as described in Jain et al ^11^. Human DNA for exon enrichment or gene targeting was sheared to approximately 12kb using g-TUBE (Covaris). All sequencing reported here was carried out on a GridION Mk1 (ONT). Standard scripts for sequencing were used with one modification, namely that the size of data chunk delivered by MinKNOW was reduced from 1 second to 0.4 seconds by changing the value of the break_reads_after_seconds parameter in the relevant TOML file (located in../minknow/conf/package/sequencing/ for MinKNOW core version 3.6). All sequencing used FLO-MIN106 R9.4.1 flowcells.

### Structural Variant Detection

Reads from NB4 cells were filtered to remove those less than 750 bases using NanoFilt ^30^. For sequencing runs using NA12878 reads were not filtered. Reads were mapped to hg38 removing ALTs with minimap2 using standard settings for ONT reads ^7^. Structural variant calling was performed with svim using default settings ^26^. Variant calls were filtered with the default filter pass and non BND (Breakpoint End) structural variant types were ignored. SVs were visualised with Ribbon ^28^.

### Code availability

The ONT Read Until API is required for running Read Until. This available from ONT. We have made minor changes to this API available from our GitHub repo. These changes are required to run in Python3 and also change the behaviour of the read cache enabling consecutive chunks of data to be stored for calling. As the ONT tool chain matures to Python3 such changes will no longer be required and these tools will be able to be run within the MinKNOW Python environment directly. PyGuppy, a python interface to the Guppy Base Calling Server is currently available on request from ONT and we hope will soon be available via PyPI. Our code is available open source at (http://www.github.com/LosseLab/ru).

### Read Until Implementation

**Table 1.**
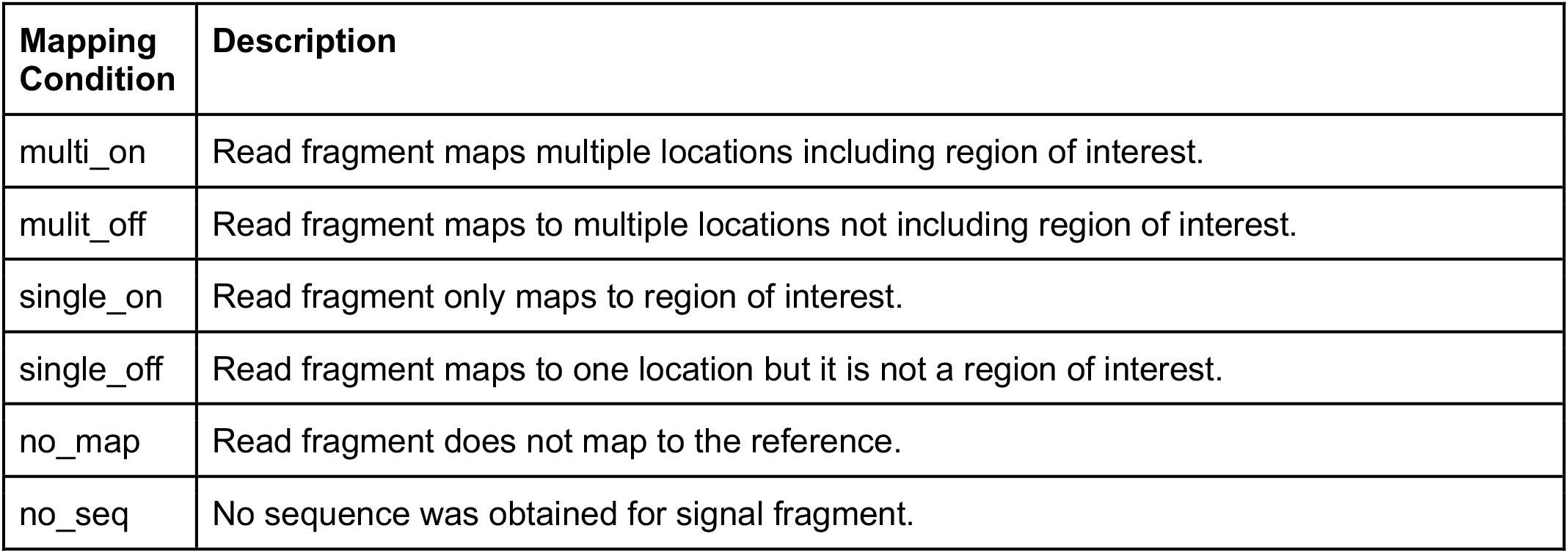
Description of possible read mapping conditions.

**Table 2.**
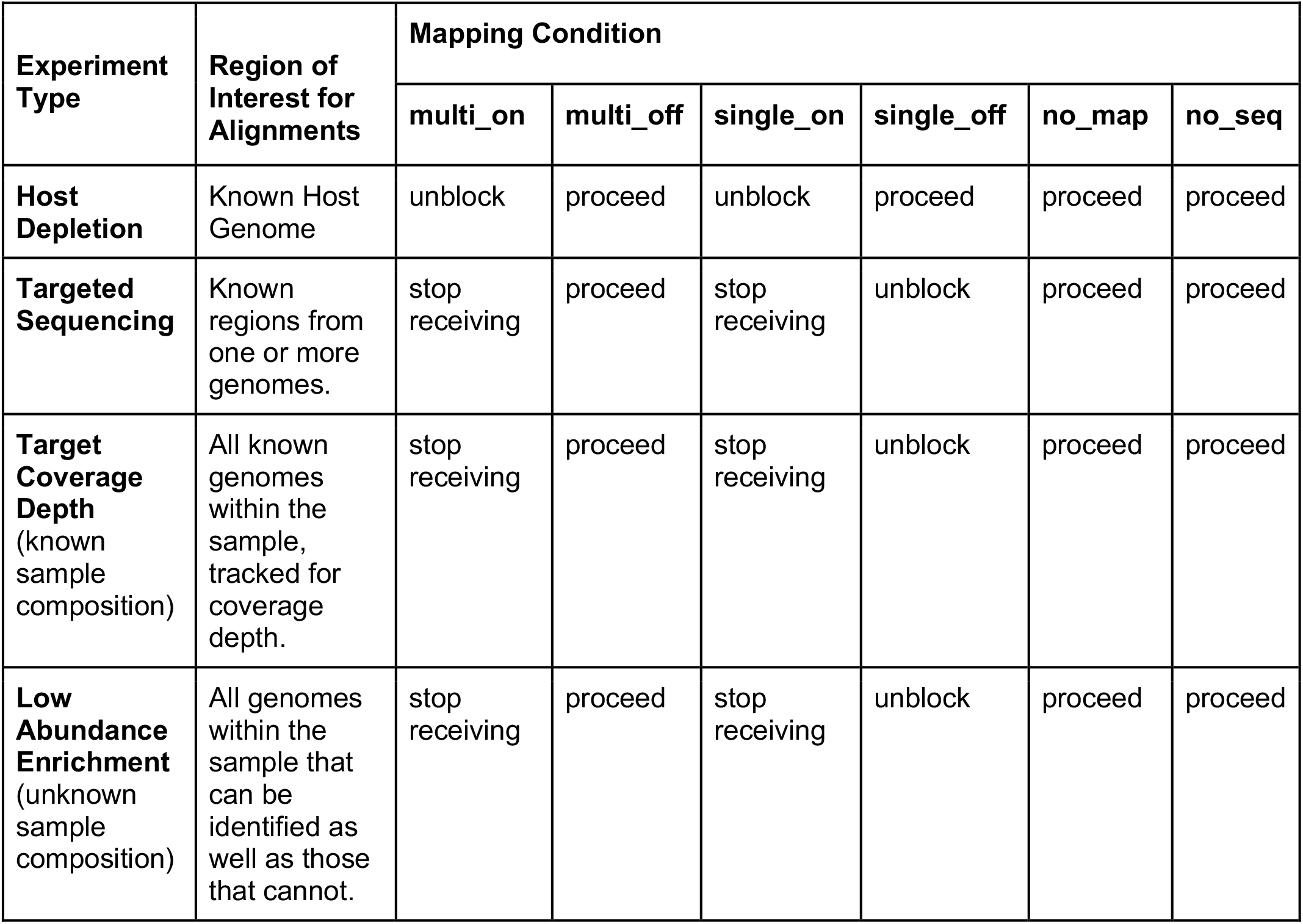
Example configurations for different experiment types. “Unblock” causes a read to be ejected from the pore, “proceed” means that a read continues to sequence and serve data through the API for later decisions, “stop receiving” allows the read to continue sequencing with no further data served through the API.

### Read Until scripts

#### ru_generators

This script runs the core Read Until process as specified in the experiment TOML file. It can select for specific regions of a genome, mapping reads in real time using minimap2 and rejecting reads appropriately. This script should be started once the initial mux scan has completed. The experiment TOML file can be updated during a sequencing run to change the configuration of the Read Until process. It is through this mechanism that ru_iteralign and ru_iteralign_centrifuge can change Read Until behaviour during a run. Configuration parameters are available under the help flag for the script. Table 1 and 2 describe the mapping parameters and configuration options for various possible experiment types.

#### ru_iteralign

This script runs an instance of the “Run Until” monitoring system that watches as completed reads are written to disk. When new data is detected ru_iteralign will map the data against the target reference genome (specified in the experiment TOML file) and compute the cumulative coverage for the sequencing run. Once a genomic target reaches sufficient coverage, it will be added to the unblock list. Optionally, the user can provide additional targets from the start of the run to implement “host depletion”. Finally, the user can configure ru_iteralign to stop the entire run if all samples have reached the required coverage depth. At present, this coverage depth is uniform for all samples, so it is not possible to have variable coverage over a target set.

#### ru_iteralign_centrifuge

This script runs an instance of the “Run Until” monitoring system. As completed reads are written to disk ru_iteralign_centrifuge will classify the reads using centrifuge and a user defined index. When 2000 reads are uniquely classified the corresponding reference genome is downloaded from RefSeq and incorporated into a minimap2 index. At this point the same process as in ru_iteralign is used to determine coverage depth. The new alignment index is passed to the core Read Until script (ru_generators) by updating the experiment TOML file allowing dynamic updates for both the unblock list and the genomic reference.

#### ru_unblock_all

This script is provided as a test of the Read Until API where are all incoming read fragments are immediately unblocked. It allows a user to quickly determine if their MinKNOW instance is able to provide and process unblock signals at the correct rate. Users should provide a bulk FAST5 file for playback for this testing process.

#### ru_validate

This script is a standalone tool for validating an experiment TOML file. We provide a ru_schema.json (https://github.com/LooseLab/ru/blob/master/ru_toml.schema.json) that describes the required configuration format.

## Acknowledgments

The authors thank Josh Quick, John Tyson, Jared Simpson and Nick Loman for helpful comments and (mainly) criticisms. Ewan Birney, Nick Goldman and Alexander Senf for helpful insights and discussion on these approaches. We thank Mike Hubank and Lewis Gallagher for access to materials and reagents as well as general boundless enthusiasm. The authors also thank Stuart Reid, Chris Wright, Chris Seymour, Jon Pugh and George Pimm from Oxford Nanopore Technologies for advice on MinKNOW and Guppy operations as well as extensive troubleshooting!

