## Supplementary material for "Nanopore adaptive sequencing for mixed samples, whole exome capture and targeted panels": Supplmentary Figures and Tables

### Supplementary Figures

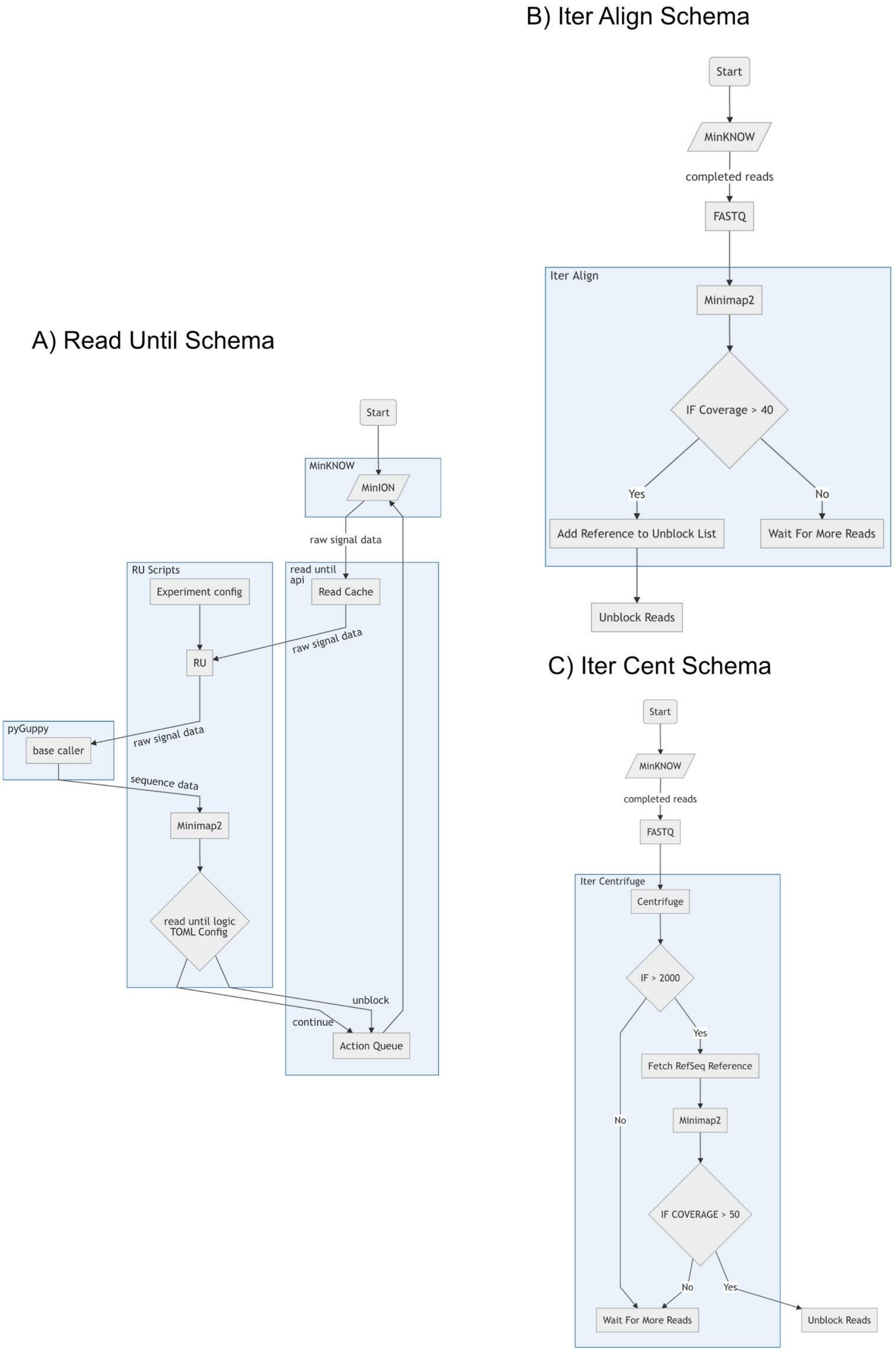


*Supplementary Figure 1: Flow diagrams of Read Until components. A) The core Read Until program which manages base calling and aligning read fragments from the device. B) The IterAlign program monitors completed reads and computes coverage based on a known reference. C) IterAlign Centrifuge monitors completed reads as in IterAlign, but uses centrifuge to classify reads and then downloads a RefSeq genome once a defined threshold is reached.*

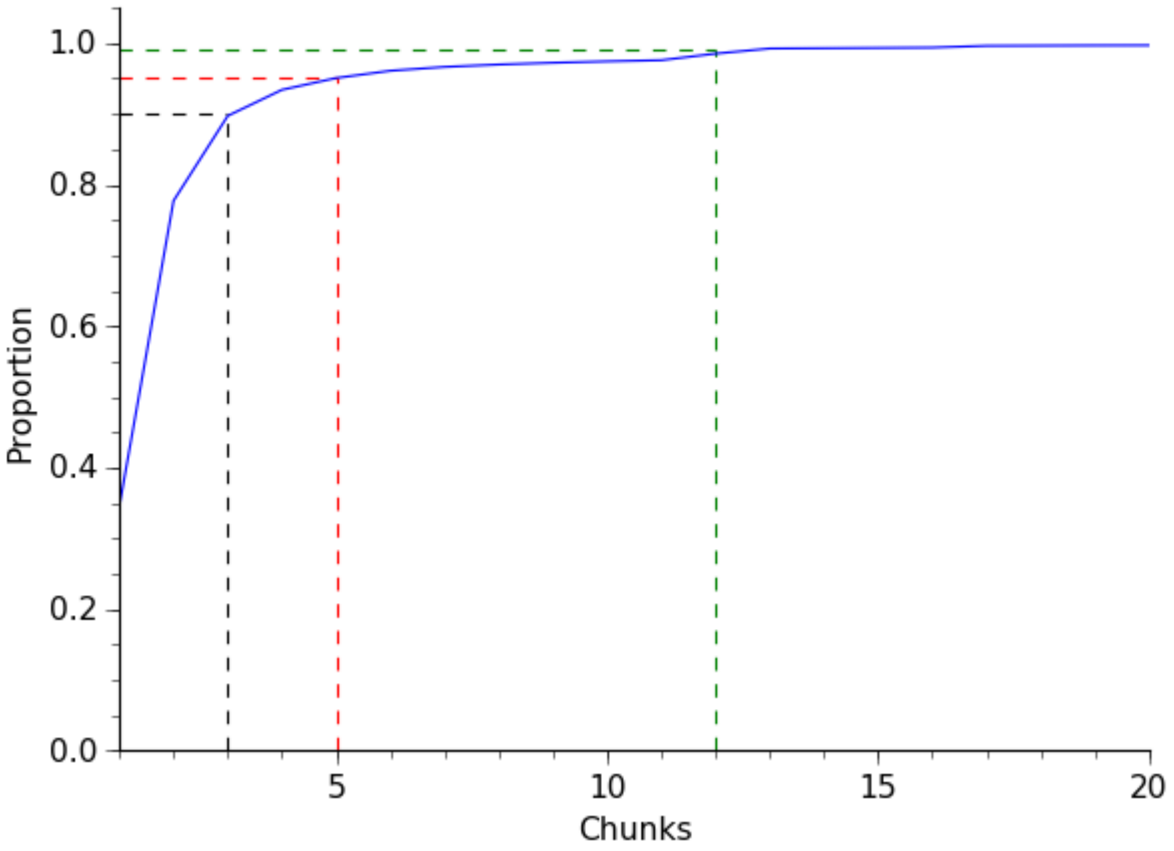


*Supplementary Figure 2: Proportion of read fragments processed in a given number of chunks. 90% of reads (black dashed line) are processed in 3 chunks, 95% (red dashed line) in 5 chunks and 99% (green dashed line) in 12 chunks.*

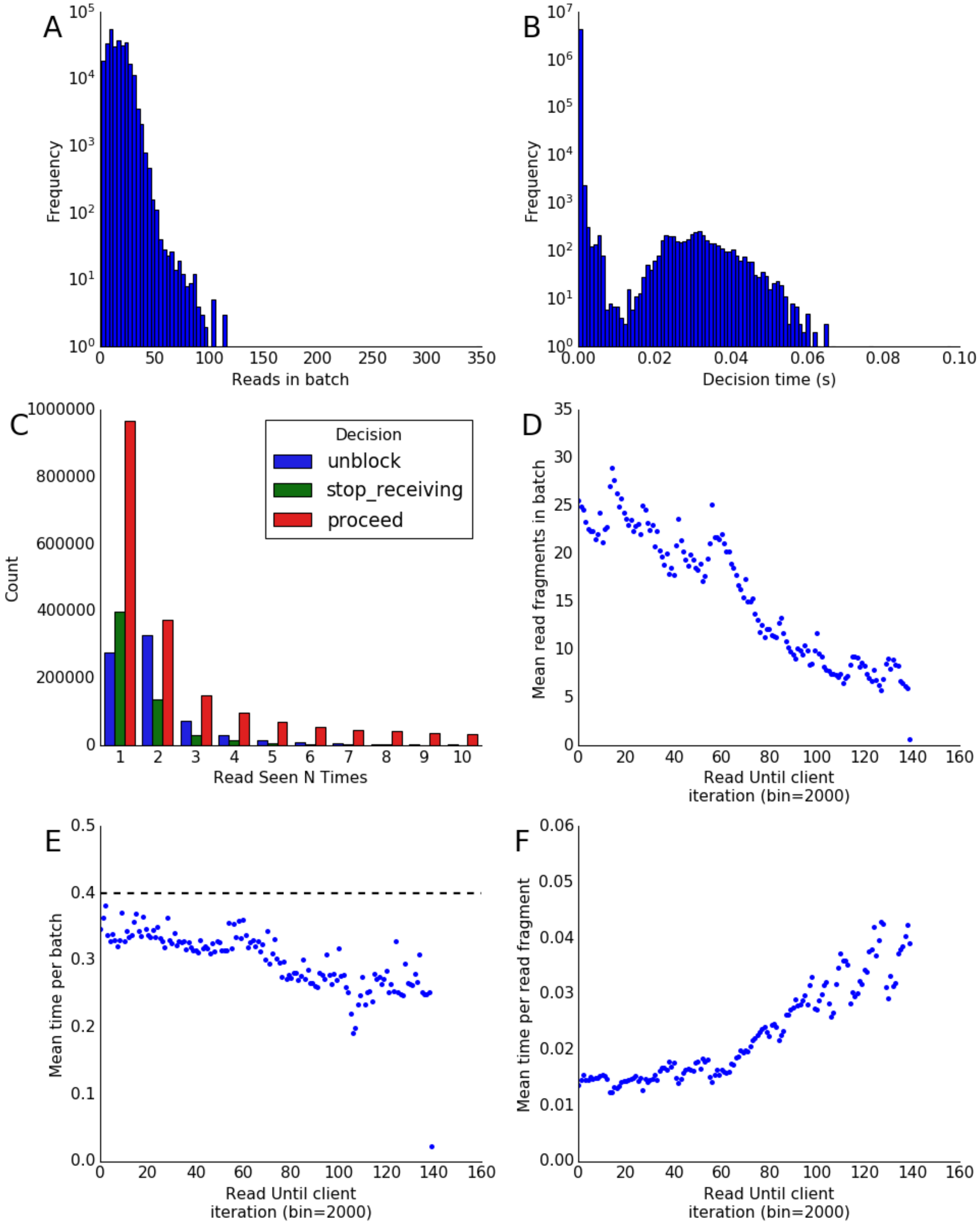


*Supplementary Figure 3: Human Chromosome Enrichment. A) Histogram of read batch size throughout the selective sequencing program. B) Histogram of decision times (time to choose unblock, stop receiving, or proceed from an alignment). C) Counts of decision classifications for read fragments seen a given number of times. D) Mean batch size, in bins of 2000, seen throughout the selective sequencing program. E) Mean process time, in bins of 2000, for batches of read fragments throughout the run. Dashed line indicates the minimum data chunk size delivered by MinKNOW. F) Mean decision time per read fragment, in bins of 2000, throughout the run. As the number of reads in a batch reduces, the overhead time of calling becomes more apparent.*

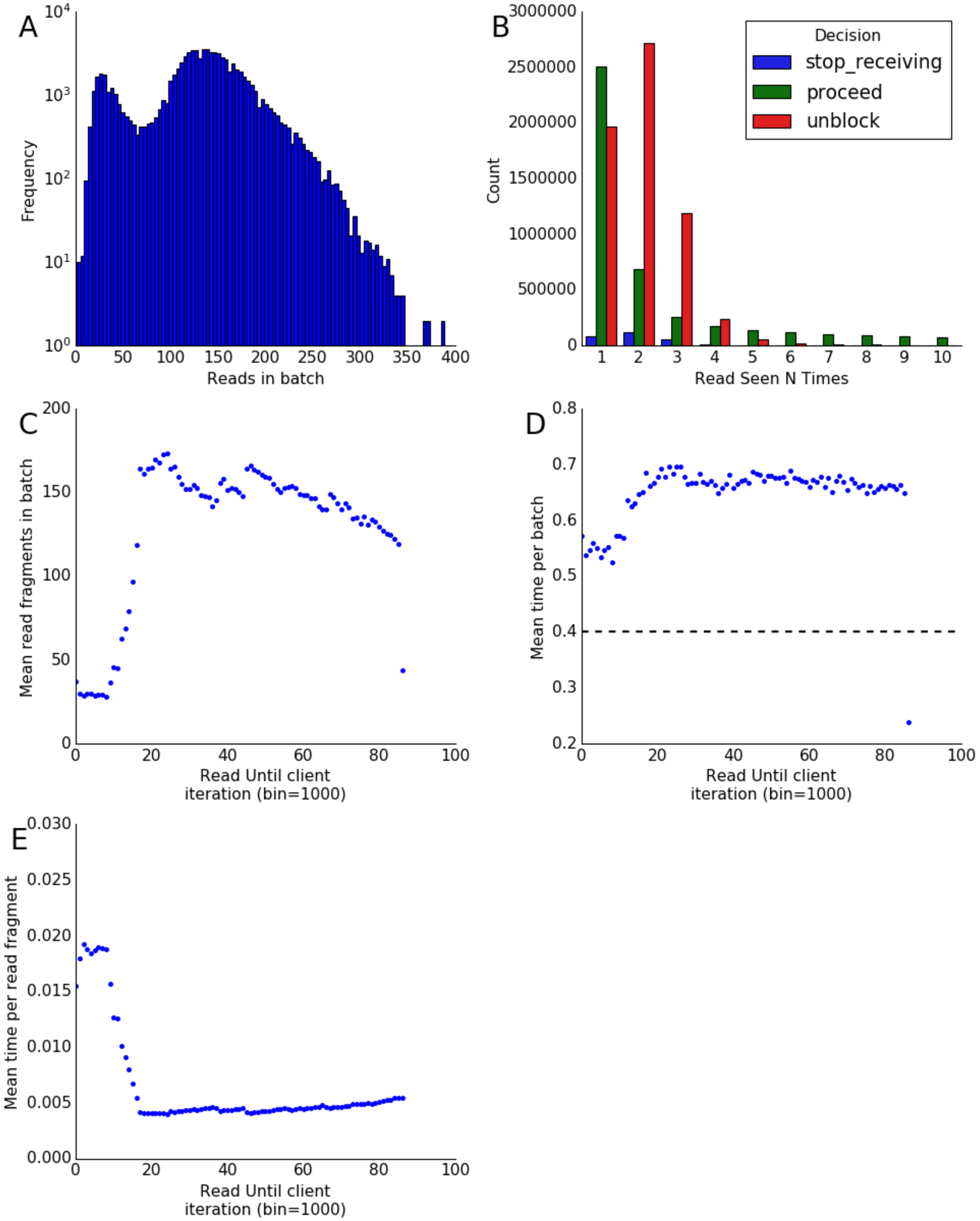


*Supplementary Figure 4: IterAlign A) Histogram of read batch size throughout the selective sequencing program. B) Histogram of decision times (time to choose unblock, stop receiving, or proceed from an alignment). C) Counts of decision classifications for read fragments seen a given number of times. D) Mean batch size, in bins of 2000, seen throughout the selective sequencing program. Dashed line indicates the minimum data chunk size delivered by MinKNOW. E) Mean process time, in bins of 2000, for batches of read fragments throughout the run. F) Mean decision time per read fragment, in bins of 2000, throughout the run. As the number of reads in a batch reduces, the overhead time of calling becomes more apparent.*

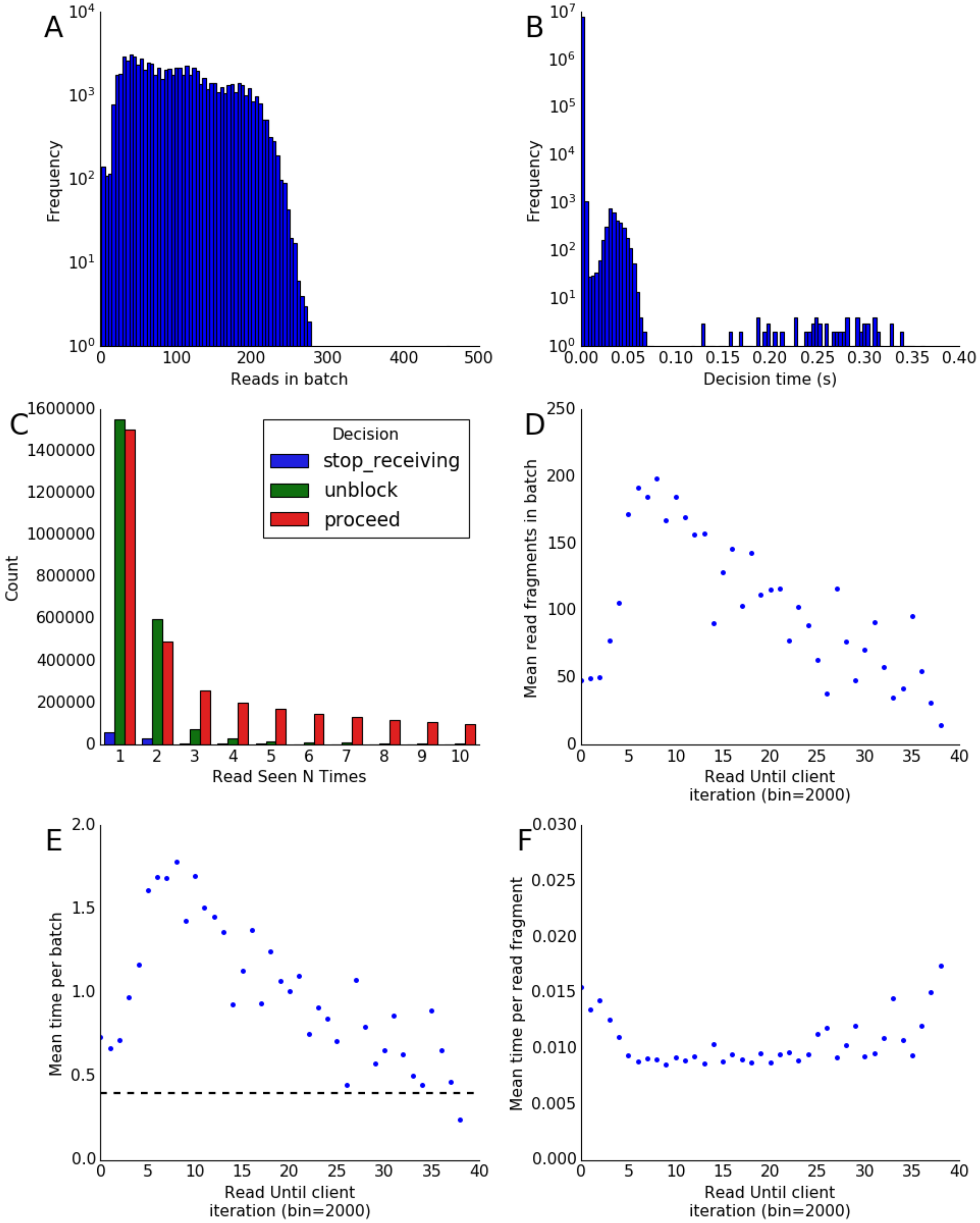


*Supplementary Figure 5: IterAlign Centrifuge Performance. A) Histogram of read batch size throughout the selective sequencing program. B) Histogram of decision times (time to choose unblock, stop receiving, or proceed from an alignment). C) Counts of decision classifications for read fragments seen a given number of times. D) Mean batch size, in bins of 2000, seen throughout the selective sequencing program. E) Mean process time, in bins of 2000, for batches of read fragments throughout the run. Dashed line indicates the minimum data chunk size delivered by MinKNOW, the longer processing times here, reflect the addition of centrifuge to the pipeline. F) Mean decision time per read fragment, in bins of 2000, throughout the run. As the number of reads in a batch reduces, the overhead time of calling becomes more apparent.*

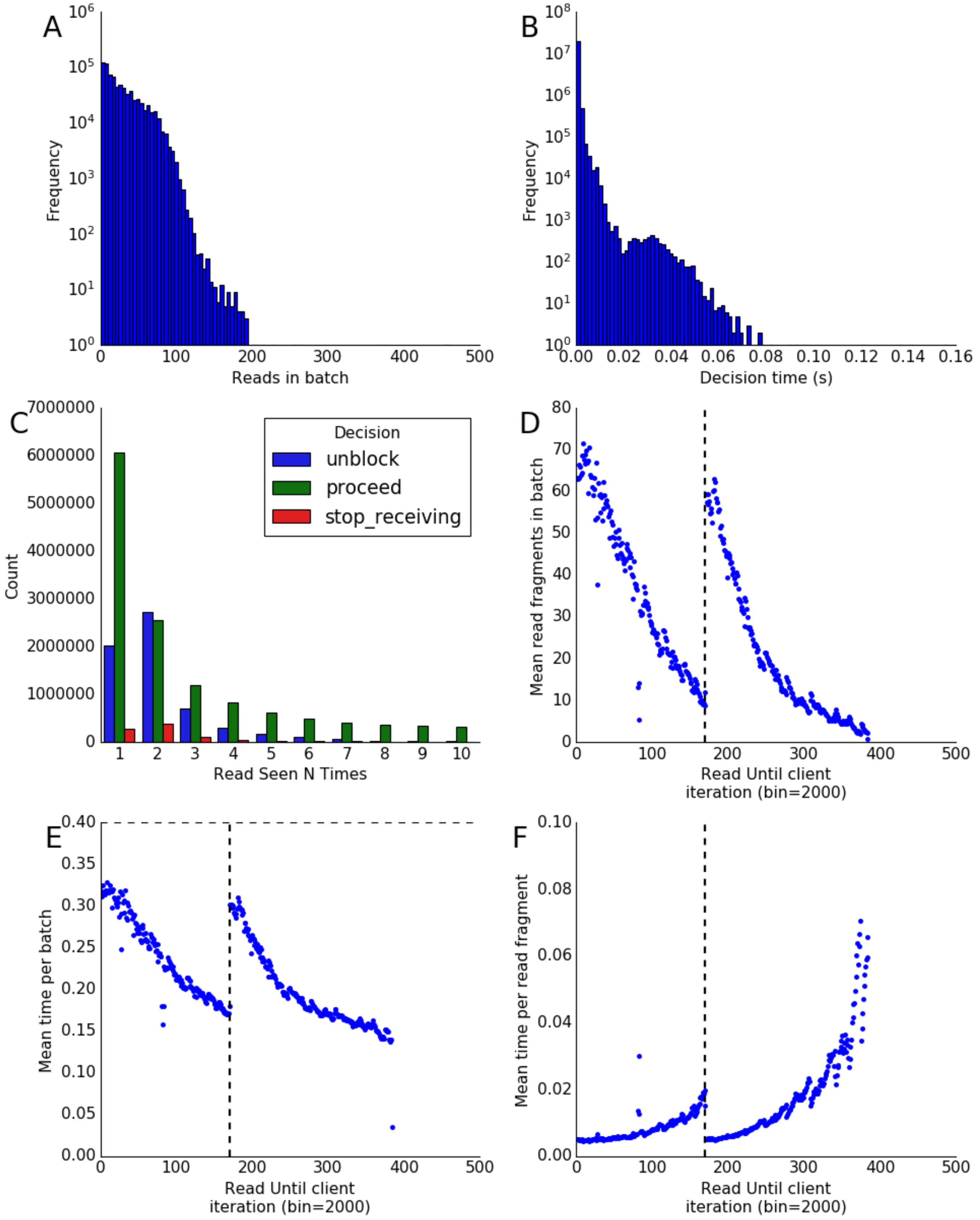


*Supplementary Figure 6: Human Exome A) Histogram of read batch size throughout the selective sequencing program. B) Histogram of decision times (time to choose unblock, stop receiving, or proceed from an alignment). C) Counts of decision classifications for read fragments seen a given number of times. D) Mean batch size, in bins of 2000, seen throughout the selective sequencing program. E) Mean process time, in bins of 2000, for batches of read fragments throughout the run. F) Mean decision time per read fragment, in bins of 2000, throughout the run. As the number of reads in a batch reduces, the overhead time of calling becomes more apparent. The vertical dashed lines mark the*

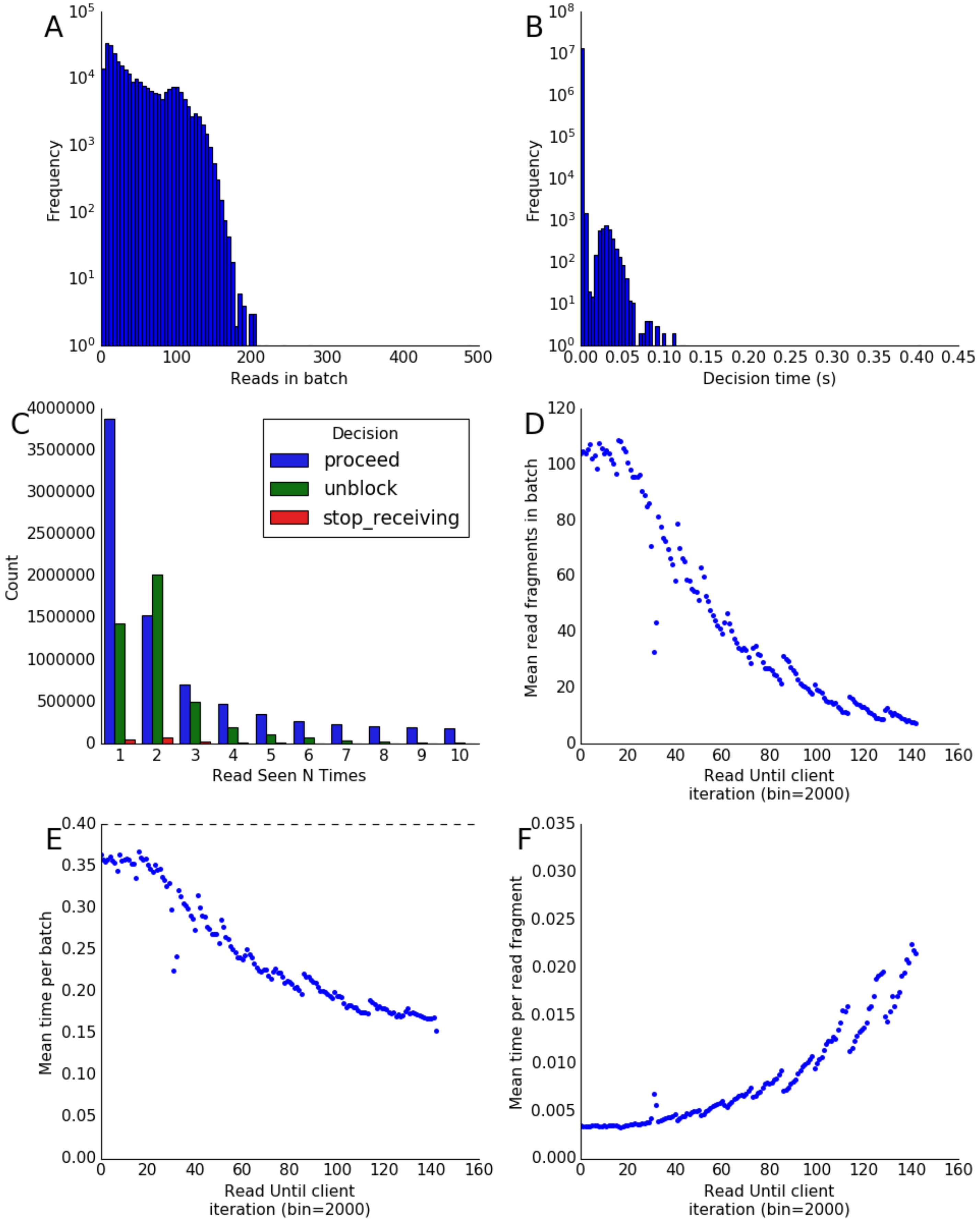


*Supplementary Figure 7: COSMIC panel A) Histogram of read batch size throughout the selective sequencing program. B) Histogram of decision times (time to choose unblock, stop receiving, or proceed from an alignment). C) Counts of decision classifications for read fragments seen a given number of times. D) Mean batch size, in bins of 2000, seen throughout the selective sequencing program. E) Mean process time, in bins of 2000, for batches of read fragments throughout the run. F) Mean decision time per read fragment, in bins of 2000, throughout the run. As the number of reads in a batch reduces, the overhead time of calling becomes more apparent.*

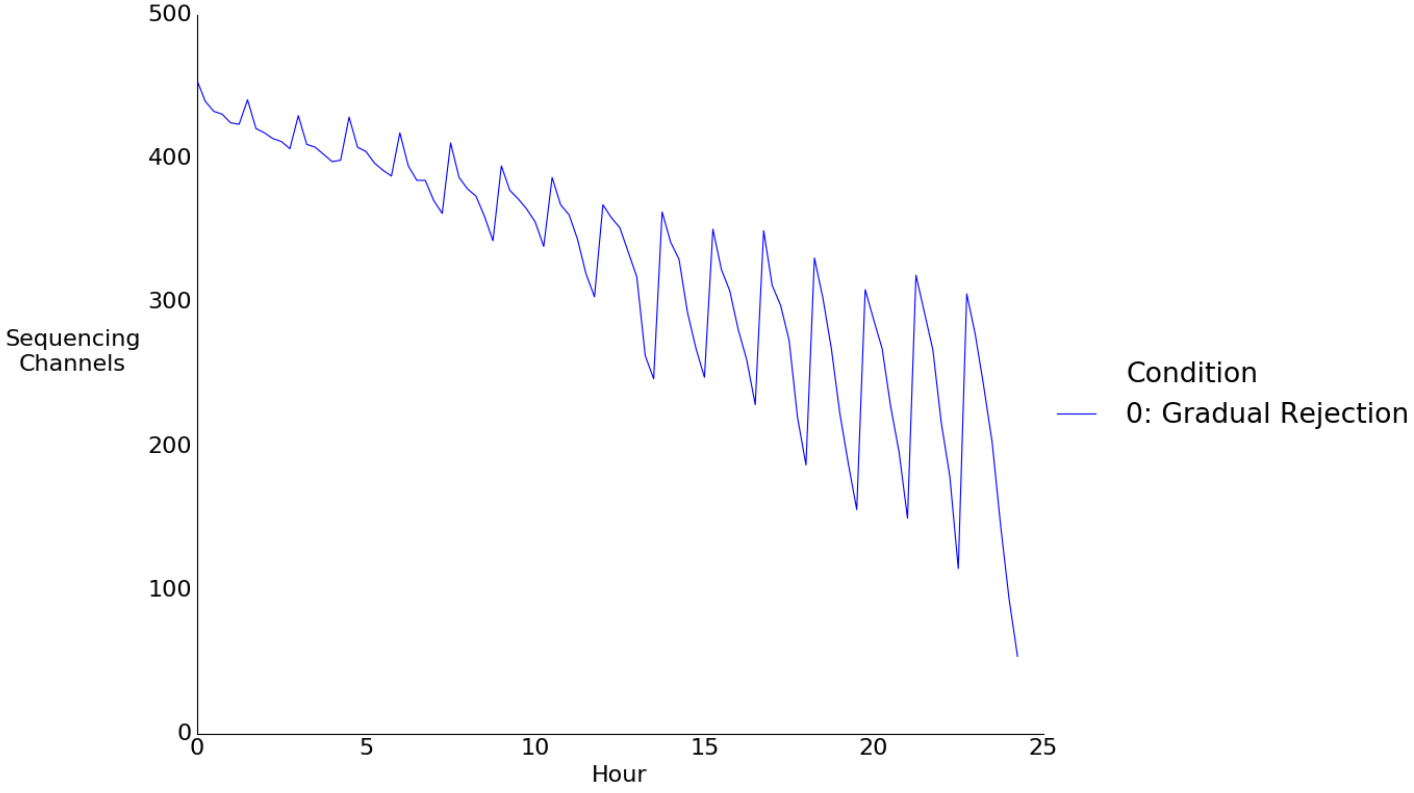


*Supplementary Figure 8: Active sequencing channels plotted over time for IterAlign Centrifuge sequencing run.*

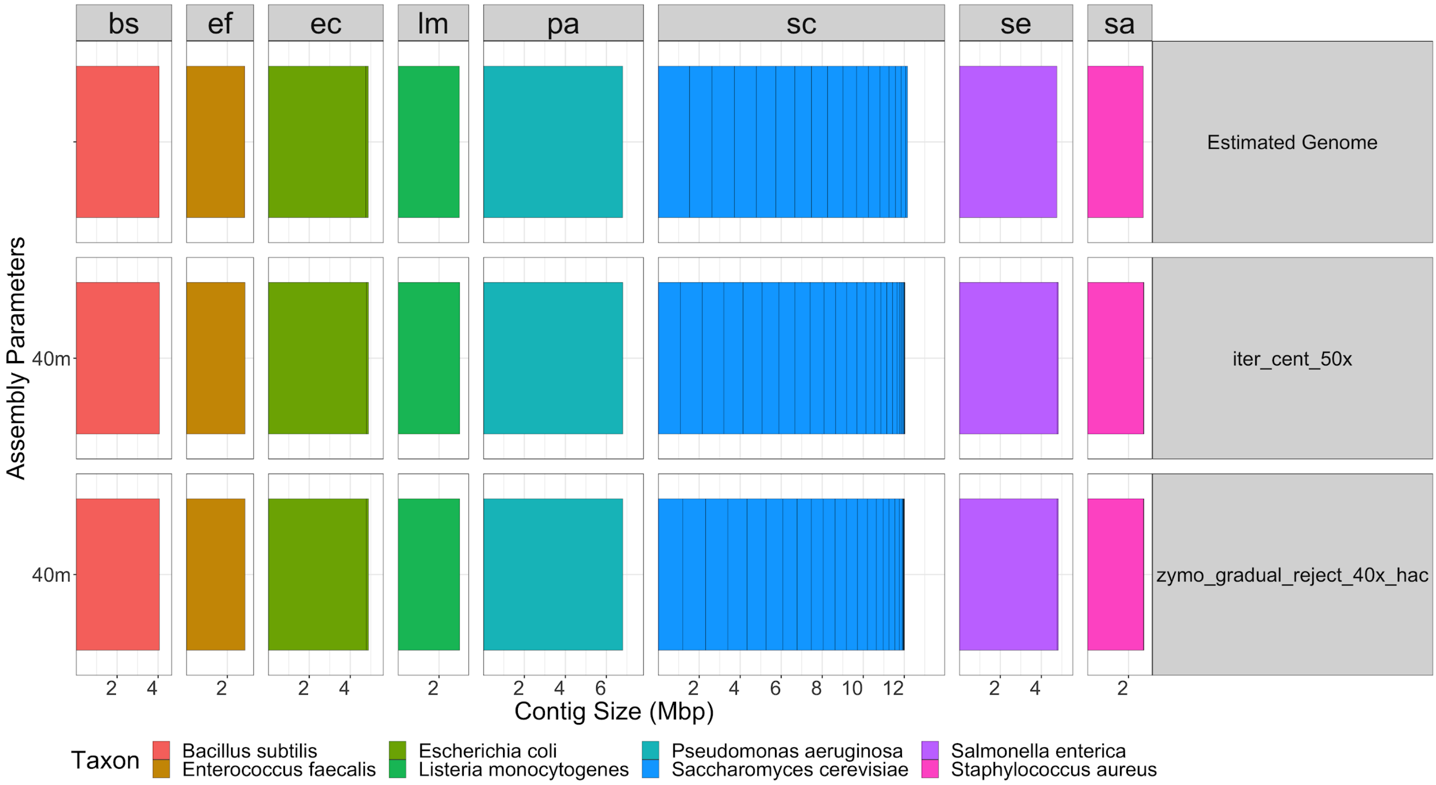


*Supplementary Figure 9. Assemblies for ZymoBIOMICS read until enriched data. Data assembled using MetaFlye using an estimated genome size of 40 Mb.*

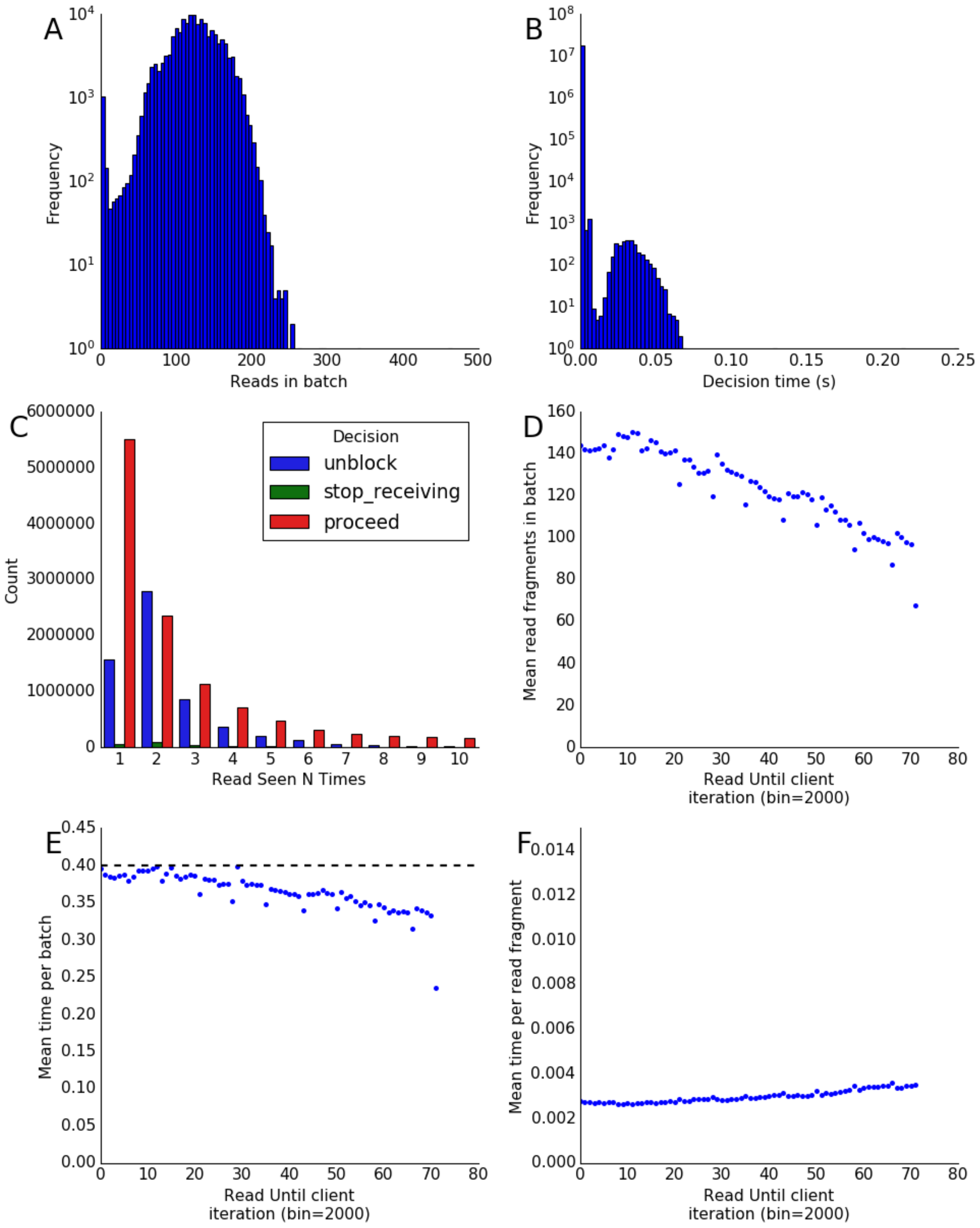


*Supplementary Figure 10: COSMIC panel against NB4 cells. A) Histogram of read batch size throughout the selective sequencing program. B) Histogram of decision times (time to choose unblock, stop receiving, or proceed from an alignment). C) Counts of decision classifications for read fragments seen a given number of times. D) Mean batch size, in bins of 2000, seen throughout the selective sequencing program. E) Mean process time, in bins of 2000, for batches of read fragments throughout the run. F) Mean decision time per read fragment, in bins of 2000, throughout the run. As the number of reads in a batch reduces, the overhead time of calling becomes more apparent.*

### Supplementary Tables

| Panel | Sample | Chrom | Pos | Qual | Filter | Alt | Support | STD POS1 | STD POS2 |
|---|---|---|---|---|---|---|---|---|---|
| E | NA12878 | chr1 | 16002843 | 7 | PASS | ]chr9:111642651]N | 6 | 3 | 1 |
| E | NA12878 | chr3 | 151430755 | 5 | PASS | [chr5:39787648[N | 4 | . | . |
| E | NA12878 | chr5 | 39787648 | 5 | PASS | [chr3:151430755[N | 4 | . | . |
| E | NA12878 | chr6 | 382460 | 7 | PASS | N]chr16:33626062] | 6 | . | . |
| E | NA12878 | chr7 | 26213350 | 6 | PASS | N]chr15:40561994] | 5 | . | . |
| E | NA12878 | chr9 | 42900334 | 6 | PASS | ]chr9:64124004]N | 5 | 5 | 19 |
| E | NA12878 | chr9 | 64124004 | 6 | PASS | N[chr9:42900334[ | 5 | 19 | 5 |
| E | NA12878 | chr9 | 111642651 | 7 | PASS | N[chr1:16002843[ | 6 | 1 | 3 |
| E | NA12878 | chr15 | 40561994 | 6 | PASS | N]chr7:26213350] | 5 | . | . |
| E | NA12878 | chr16 | 33626062 | 7 | PASS | N]chr6:382460] | 6 | . | . |
| C | NA12878 | No BND variants detected in the COSMIC Panel for NA12878. | | | | | | | |
| C | NB4 | chr3 | 136559276 | 7 | PASS | N[chr3:136714904[ | 6 | . | 9 |
| C | NB4 | chr3 | 136714904 | 7 | PASS | ]chr3:136559276]N | 6 | 9 | . |
| C | NB4 | chr15 | 74034025 | 7 | PASS | ]chr17:40345927]N | 6 | 2 | 5 |
| C | NB4 | chr17 | 40345927 | 7 | PASS | N[chr15:74034025[ | 6 | 5 | 2 |

*Supplementary Table 1: Structural variants as identified by svim. Variant calls were filtered with the default filter pass and non BND (Breakpoint End) structural variant types are not shown in this table. Panel: E - Exon Panel, C - COSMIC Cancer Panel, Sample: NA12878 reference cells or NB4 reference cells. Chrom: Starting chromosome of the read. POS: Position of the break point with respect to chromosome in Chrom. Qual: Quality as reported by SVIM. Filter: Default SVIM filter (>Q5 ). Alt: Description of the second break point in the structural variant. Support: Number of reads supporting the observed structural variant. STD POS 1/2: The standard deviation in position of the breakpoint.*

#### Supplementary Files

*Supplementary File 1 - COSMIC_Coverage.csv - File containing coordinates used for selective sequencing of the COSMIC cancer panel along with the observed mean coverage, gene name and span of the region being enriched for. All coordinates with respect to hg38.*
